# Are global hotspots of endemic richness shaped by plate tectonics?

**DOI:** 10.1101/131128

**Authors:** Pellissier Loïc, Christian Heine, Camille Albouy

## Abstract

Singular regions of the globe harbour a disproportionally large fraction of extant biodiversity. Spatial biodiversity gradients are frequently associated to extant ecological conditions using statistical models, but more rarely to paleo-environmental conditions, especially beyond the Quaternary. On one hand the role of plate tectonics in shaping the extant diversity of lineages is supported by numerous phylogenetic and fossil evidences, and on the other hand the spatial variation of biodiversity across the globe is rarely associated to geodynamic variables. In this study, we propose that plate tectonics explain the current location of hotspots of endemic richness across the globe. As an illustration, we used paleogeographies in a model, which quantifies through time and for each cell the potential dispersal across disconnected habitat patches. Rare events of dispersal across dynamic straits of unsuitable habitats allows species colonisation and that a subsequent absence of gene flow could lead to in-situ speciation. We evaluated whether this process could pinpoint the locations of hotspots of endemic richness computed from the ranges of 181,603 species across 14 taxonomic groups. The significant congruence between the regions highlighted by the model and the endemic richness provides evidences of the contribution of plate tectonics in shaping global biodiversity gradients. Places with high tectonic complexity, predominantly located at the confluence of major lithospheric plates such as the Mediterranean basin, Central America, Madagascar and South East Asia likely provided favourable circumstances for allopatric speciation and the emergence of new species across straits. While our illustration supports the role of plate tectonics, accounting for deep time geological events in spatial models of extant biodiversity is not straightforward. Future research should develop quantitative spatial models of biodiversity including the dynamic of ancient habitats.

## Introduction

Biodiversity on Earth is the result of the radiation of lineages in interaction with relentless changes in their environment and is sustained by extant ecological conditions. Convergence of ecological and evolutionary theories stems from the recognition that the uneven spatial distribution of biodiversity is the product of both contemporary and historical factors (Latham and Ricklefs 1993, Mittelbach et al. 2007). Investigations of the emergence and maintenance of biodiversity has been addressed using a variety of approaches. Based on phylogenies, rates of diversification have been associated to regional differences in paleo-environmental conditions and highlighted how ancient Earth processes lead to extant biodiversity (e.g. Near et al. 2012, Condamine et al. 2012). In particular, plate tectonics were shown to foster species diversification in specific regions (Magri et al. 2007, Li et al. 2013, Richardson et al. 2014, Bagley and Johnson 2014). However, phylogeographic models are based on a very limited number of regions (e.g. low versus high latitude, Pyron 2014, Pulido-Santacruz and Weir 2016), and their power to explain complex spatial gradients is limited. Alternatively, spatial statistical models provide a popular approach to explain spatial gradients of biodiversity. In spatial statistical models, contemporary factors are more frequently investigated (e.g. climate or energy, Currie 1991, Hawkins et al. 2003, Jetz and Kreft 2007), than historical proxies (e.g. latitude, Kerkhoff et al. 2014), which rarely extend beyond the Quaternary (Graham et al. 2006, Sandel et al. 2011, Pellissier et al. 2014). While plate tectonics are recognised to have shaped past and extant biogeographic patterns (Valentine and Moores 1970, Valentine 1971, Raven and Axelrod 1975, Cracraft 1973, Flessa 1980, Briggs 2003), proxies of ancient geological dynamics are not integrated in spatial models of biodiversity. Hence, correlative climatic models may currently underestimate the influence of historical factors on extant biodiversity patterns (Rahbek et al. 2007).

Using spatial statistical models, the large scale distribution of biodiversity across the globe has been associated to a multitude of variables (Mittelbach et al. 2007). The most commonly proposed driver of biodiversity is energy, either in the form of temperature (Currie 1991, Hawkins et al. 2003) or primary productivity (Kay et al. 1997, Waide et al. 1999, Jetz and Fine 2012), where higher energy should sustain more complex food chains (Briand and Cohen 1987). Other extant environmental conditions, including water (Qian et al. 2007) or area (Hawkins et al. 2003, Bellwood et al. 2005) show strong correlations with extant species diversity. Most historical studies investigate whether species distributions are in disequilibrium with extant climate as a legacy of the last glaciations (Graham et al. 2006, Sandel et al. 2011, Pellissier et al. 2014), while few spatial analyses extend beyond the Quaternary (Svenning et al. 2015). Yet, phylogeography and palaeontological evidences suggest that geological dynamics largely reshaped species distribution in the past (Li et al. 2013, Richardson et al. 2014, Bagley and Johnson 2014), and their legacy should be detectable today. Moreover, the association between species richness and topography (Davies et al. 2007), heterogeneity (Stein et al. 2014) or plate boundaries (Keith et al. 2013) suggests that older processes, e.g. associated to orogeny, also represented biodiversity pumps (Badgley 2010). Models of the spatial distribution of extant biodiversity should integrate proxies of more ancient processes, which acted concomitantly to the phases of diversification of extant species, usually millions of years old (Hodge and Bellwood 2016).

Beyond a single latitudinal gradient, singular regions of the globe host a disproportionally large fraction of extant biodiversity. For instance, at comparable latitudes and climates, Asia is considerably more diverse than America or Africa (Ricklefs et al. 2004, Couvreur 2015). During the last ∼ 100 million years, a period during which many extant clades diversified, plate tectonics largely remodelled the structure of terrestrial landscapes and marine reefs and should have left a legacy on current diversity (Renema et al. 2008, Leprieur et al. 2016). Among geological processes shaping biodiversity, the convergence of tectonic plates promotes environmental heterogeneity through building of topography. Here, temporal changes in convergence kinematics along subduction zones (such as the Andes) can generate significant vertical crustal motions. Towards the end of a Wilson cycle, collision of continental plates generate Alpine-Himalayan type mountain chains, which enhance the diversification of organisms, e.g. plant (Hoorn et al. 2010) and animal lineages (Toussaint et al. 2014). Additional geological processes, e.g. the emergence of islands associated to volcanism or the collision of continental plates, modulated species diversification (Briggs 2003, Bidegaray-Batista et al. 2011). Although orogeny and island formation processes are commonly known to isolate populations and promote speciation (Hoorn et al. 2010, Ali and Aitchison 2014), those represent only part of the full spectrum of plate tectonic processes. By isolating populations with oceanic gateways, fuelled by sporadic events of long-distance dispersal across straits (Lavergne et al. 2013), plate divergence may be a major catalyst of allopatric speciation (Steeman et al. 2009). The full spectrum of relative plate motions, including the plate divergence and the resulting formation of oceanic gateways and basins, had probably the most profound effect on the diversification of species through allopatric speciation and may have led to biodiversity unbalance across the globe.

In this article, our aim is to highlight that, despite phylogeographic and fossil evidences, most recent studies seeking to understand the cause of the spatial variation of extant biodiversity using spatial model have overlooked plate tectonics. Ancient geological events are expected to have promoted differential diversification across the globe fostering global imbalance in biodiversity (Gillespie and Roderick 2014). We illustrate our hypothesis using a simple spatial model which quantifies dynamically the role of plate movements in providing opportunities to generate biodiversity at the margins of continental plates. Allopatry is thought to be the dominant force of speciation (Futuyma and Mayer 1980) and this mechanism may generate a higher diversity of species endemic to regions with singular geological dynamics. We compared locations where high diversity is expected from a speciation process linked to plate movements to the distribution of the cells with the highest (top 10%) endemic richness worldwide across 14 taxonomic groups in marine, freshwater and terrestrial environment amounting to 181,603 species. Based on those evidences, we argue that the availability of paleo-habitat reconstructions should provide the missing link between past environmental changes and their consequence on present biodiversity gradients. We stress the need for further research into the role of past habitat dynamic associated to plate tectonics on the spatial distribution of current biodiversity.

## A spatial proxy of speciation from plate tectonics

Reconstructions of paleo-habitats is needed to understand which regions were historically the most favourable to promote the emergence of biodiversity (Svenning et al. 2015). Here, we used paleo-reconstructions of the absolute position of continents, coastlines and paleobathymetry (Müller 2008, Heine et al. 2015) from the Early Cretaceous (140 Ma) to the present in 1 Ma steps as boundary conditions for our models. The period spanning the last 140 Ma is characterized by significant changes in the position of the tectonic plates across the globe, e.g. from the breaking of Gondwana supercontinent, followed by the rapid northward motion of India, and Australia. This phase represents the time window during which many extant clades diversified (Meredith et al. 2011, Jetz et al. 2012). Yet, because the peak of diversification of some of the groups analysed occurred later, we run a sensitivity analysis, which showed that considering different starting dates for the simulations (100, 80, 60 and 40 Ma) provide similar distributions of expected diversity (Supplementary Appendix Fig. S1).

To produce a spatial proxy of how plate tectonics might have shaped spatial biodiversity gradients, we used paleo-environmental maps to quantify through time and for each cell the potential dispersal from disconnected land patches (Figure 1, Supplementary Appendix Fig. S1, S2). Dispersal across geographic barriers should allow the establishment of new species, but that gene flow is subsequently almost inexistent leading to in-situ speciation. Classical example of long distance colonisation of remote areas such as islands(e.g. Guzmán and Vargas 2009, Gillespie and Roderick 2014) leading to local speciation support the possibility of dispersal across straits on geological time scales (Cowie and Holland 2006). For terrestrial and freshwater organisms, land was considered as suitable and sea was considered unsuitable, while for marine reef organisms, shallow reef habitat was considered as suitable, while deeper waters were considered as unsuitable. We tested a range of dispersal *d* (5, 10, 15, 20, 25, 30 [°]) for both terrestrial and marine ecosystems (Supplementary Appendix, Fig. S4, S6). The proxy only covers a mechanism for the generation of new species, but fails to account for extinction. Extinction is supposed to be associated to climatic fluctuation more intense toward the poles (Dynesius and Jansson 2000). Hence, we also filtered the output maps with an extinction filter that decreases linearly or with a Gaussian shape with latitude (Supplementary Appendix, Fig. S5, S7).

**Figure 1:**
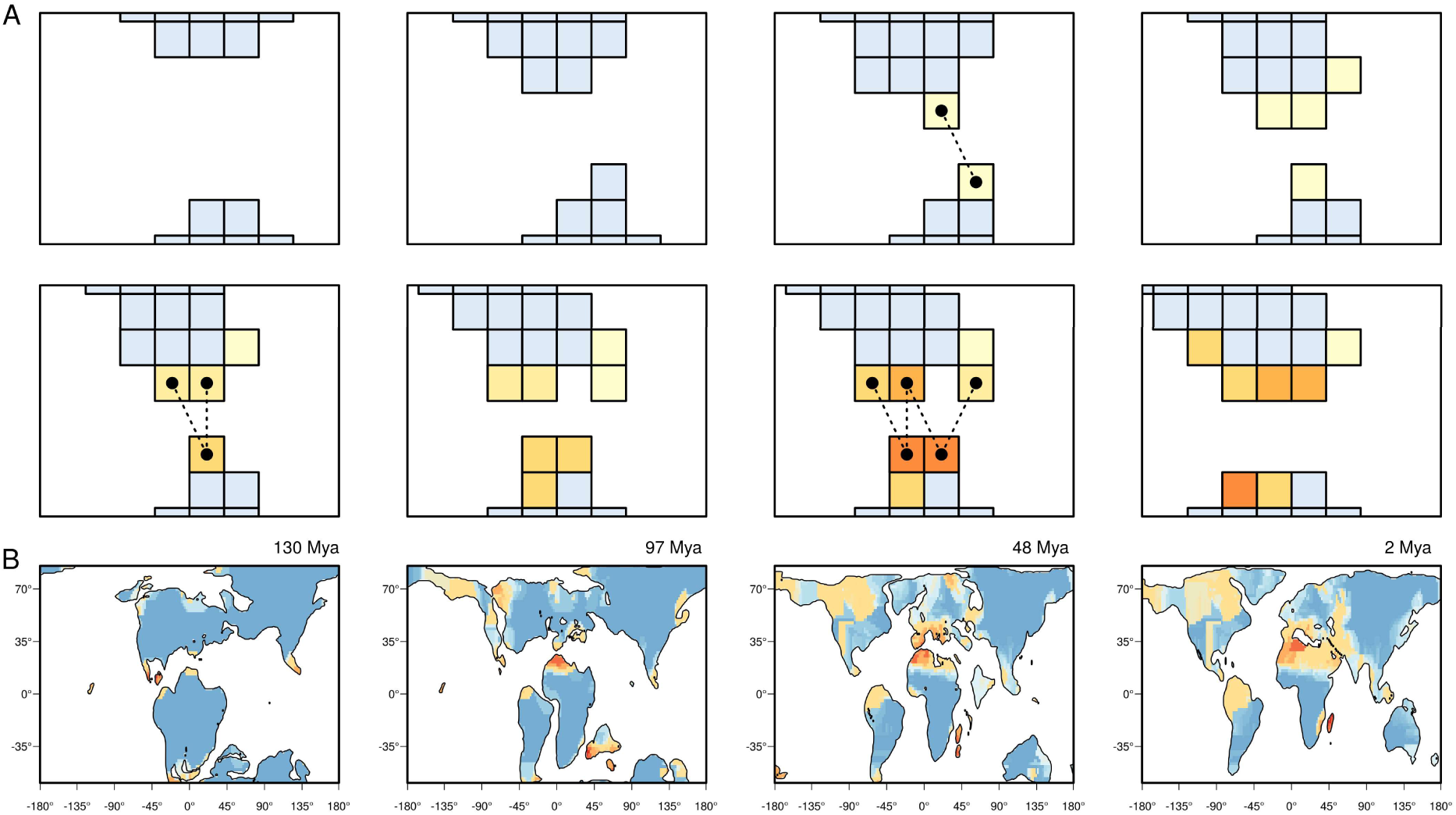
A. Illustration of the mechanism used to produce the spatial proxy of speciation from plate tectonics. During each million-year time slice, the simulation first defines the patches separated by at least one cell of unsuitable habitat using an algorithm of cluster splits implemented in the “dbscan” function in the fpc library in R. The model identifies for each cell, the number of surrounding “donor” cells from disconnected habitat patches below a dispersal distance *d*. A higher number of possible donor cells is assumed to increase the chance of long distance dispersal across a barrier. To accommodate the movement of the continents, the cell values at time t are transferred to the closest new cell values at time t+1. B. Examples of four paleogeographies showing the location of high potential for speciation across straits for 130, 97, 48 and 2 Mya.

**Figure 2:**
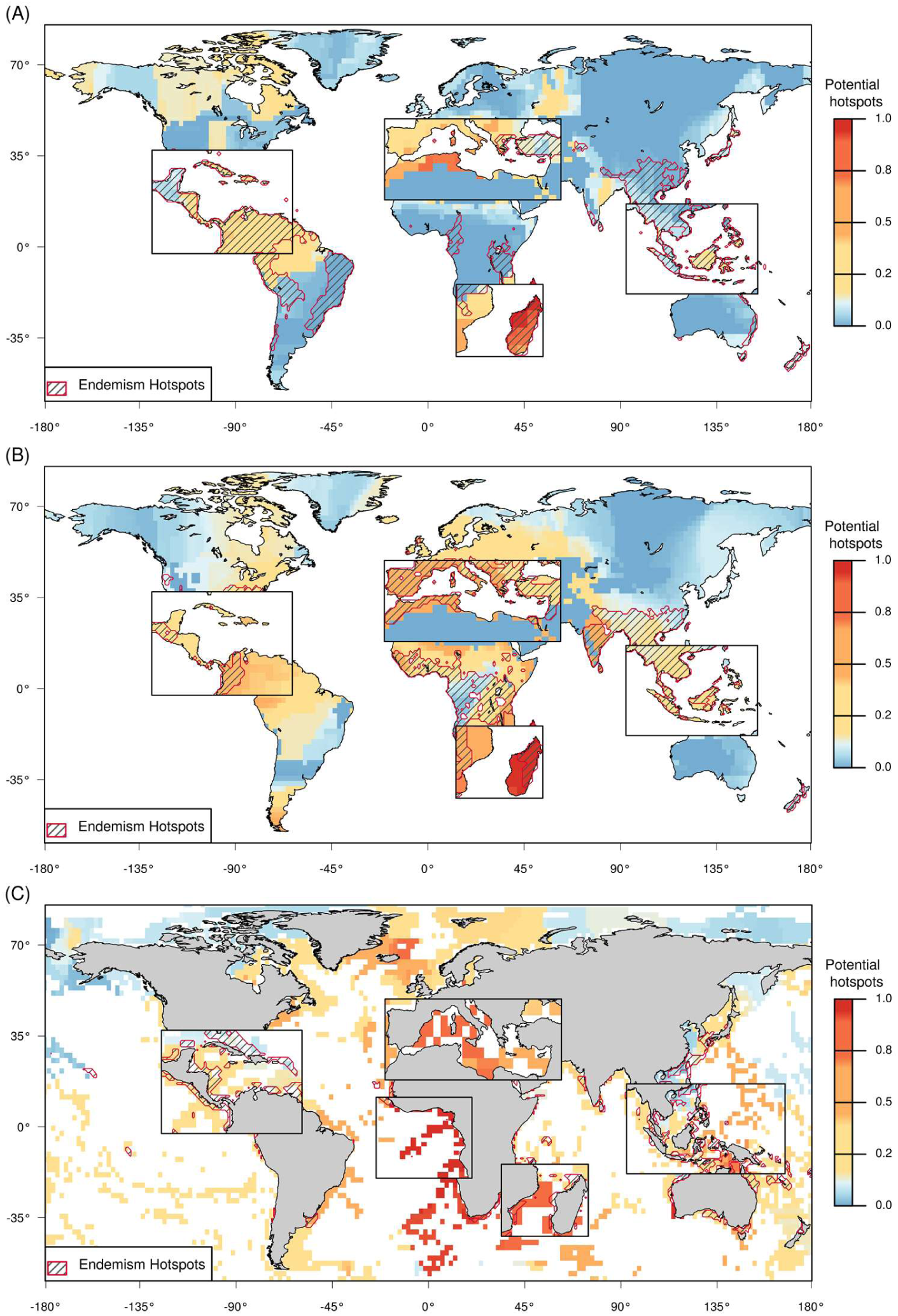
Map of the proxy of the potential for speciation associated to plate tectonics for the (A) terrestrial, (B) freshwater (C) marine species (d = 10 for A and C and d = 25 for B). The distribution of cells above the 10% of the highest values of endemic richness are represented by a smoothed hatched shape. The most important areas which provided favourable circumstances for speciation are highlighted by the zoom. Those correspond, for the terrestrial taxa, to the Mediterranean basin, Central America, Madagascar and South East Asia.

We used a global database on the geographical distribution of species obtained from BirdLife and the IUCN (IUCN Global Red List Assessments), the Ocean Biogeographic Information System and the Kew plant regional checklists. We transformed individual species shapefiles into equal area raster grids at a resolution of 1° (i.e. ∼ 111.195 km; see Fig. S8, S9, S10 in Appendix S1). We calculated the endemic richness as the sum of the inverse range sizes of all species for each taxonomic groups (Kier et al. 2009). We tested whether there is a significant spatial congruence between the singularities expected by the model and the observed endemic richness. For each comparison between observed and predicted maps, we evaluated the spatial match between the 10% of the cells with the highest endemic richness and the 10% with the highest expected value predicted by the model. To assess whether the observed number of overlapping cells is significantly different from that obtained by chance, we used a randomization procedure as in Mouillot et al. (2011). We randomly redistributed the value of expected diversity 999 times within the grid. The observed numbers of overlaps were then compared to their respective null distributions to calculate p-values. This analysis allows identifying if expected and observed diversities present similar spatial repartition of high values. This approach is more appropriate to evaluate our spatial hypothesis, but the results are also supported by standard correlations (Table S1, S2).

The resulting proxy of speciation associated to plate tectonics pinpoints a few regions with high potential for speciation associated to plate tectonics, namely Central America, the circum-Mediterranean basin, Madagascar and the Sundaland region of South East Asia. Two additional regions at higher latitude in the archipelago of Northern Canada and between Patagonia and Antarctica are also highlighted, but their general influence is attenuated once the extinction is considered (Supplementary Appendix, Fig. S11, S12). The regions which have undergone significant tectonic deformation potentially generating species across straits showed a significant match with the hotspots of endemic richness, with 28.79% of common grid cells (47.19% when considering extinction) for land organisms and 26.94% (25.41% with extinction) for freshwater organisms. Plate tectonics generated singular spatial configurations of land masses, which possibly favoured the emergence of species with restricted ranges at the land margins, hence well-matching endemic richness. Applied to the marine realm, the model pinpoints broader regions, including the Indo-Australian archipelago, the region of Madagascar, the circum-Mediterranean Sea, and the Southern Atlantic. The congruence of the model expectations and the regions of singular endemic richness was low with only 11.65% of match (24.71% with extinction).

The dynamics of continental separation and the formation of narrow, deep water oceanic basins likely permitted long-distance dispersal followed by allopatric speciation (Xiang and Soltis 2001, Lavergne et al. 2013) and corroborates the idea that plate tectonic processes over time and space shaped regions of exceptional endemic richness (Gillespie and Roderick 2014, Couvreur 2015). For instance, phylogeographic inferences suggest that dispersal from eastern Asia to insular areas in South East Asia coincides with the timing of major tectonic activity and the collision between the Sunda and Sahul shelves that caused continental fragments to be rearranged spatially throughout the region (Bacon et al. 2013). When populations are established after dispersal across oceanic gateways, isolation prompts the emergence of disparate gene pools which ultimately leads to speciation (Ali and Aitchison 2014). The model was able to highlight some of the regions of the globe with the highest level of endemic richness including the Mediterranean basin, Meso-America, Madagascar and South-East Asia (Figure 3 A). In agreement with our model, evidences suggest that periodic dispersal events across land masses triggered species diversification, with known examples being those between the North, Central and South American plates (Bagley and Johnson 2014), across the Iberian, Adriatic and African tectonic plates in the Mediterranean basin (Lavergne et al. 2013), or in the Indo-Australian archipelago (Warren et al. 2010). Similarly, the early emergence of Madagascar in interaction with asymmetrical exchanges with the coast of Africa likely enhanced speciation and promoted a high level of endemism (Ali and Huber 2010). Moreover, a paleo-strait at the Cape Horn putatively shaped high diversity in the South America-Antarctic Peninsula region, which corresponds to an exceptional fossil plant diversity in the Eocene (Wilf et al. 2005). Overall, the model shows a good match with observations for the land, but a lower match for the marine realm. We hypothesize that deep water oceanic gateways between shallow shelfal areas are probably much more permeable for marine organisms than shallow epicontinental sea strait across land patches for terrestrial taxa.

## Multi-temporal spatial maps of paleo-environments

The current climate constrains the number of species that can survive at a given place, for instance through processes of environmental filtering along latitude (Sommer et al. 2014). It is however not sufficient to explain the full regional variability of species diversity across the globe (Ricklefs et al. 2004, Ricklefs and He 2016). Complementing ecological factors, biodiversity gradients can be related to historical events of speciation and extinctions, whose intensity varied across regions (Svenning et al. 2015). Interestingly, many regions recognised for their high diversity or degree of endemism, such as Madagascar, or South East Asia, are situated in proximity to continental plate margins, where previously joined continental crust was rifted apart, providing clues on the role of plate tectonics in shaping biodiversity hotspots (Couvreur 2015). We highlight that, while many phylogeographic or paleontological evidences suggest an important role of plate tectonics, most spatial model of extant biodiversity have overlooked this effect. Here, we provide a coarse spatial proxy of plate tectonics on speciation for terrestrial dynamics to support our hypothesis. We acknowledge that our example has limitation, e.g. not integrating orogeny, thus failing to highlight regions where high endemic richness is linked to topographic complexity, such as the Andes. Improving the details of phylogeographic reconstructions will allow better quantifying how plate tectonics shaped the extant spatial gradients of biodiversity.

In recent years, significant progress has been made to reconstruct the time-dependent bathymetric evolution of ocean basins over the past 230 Million years (Müller et al. 2008, 2016), along with advances to assimilate global data sets on the distribution of paleoshorelines (Heine et al. 2015). Paleobathymetric reconstructions have been used to model the diversification of marine organisms (Leprieur et al. 2016). For terrestrial life, one of the key boundary conditions are the availability of robust paleoelevation models in regularly spaced intervals. While such models have been assembled for proprietary studies in industry-research consortiums, they are currently not freely available for academic research. Some have been used for instance to reconstruct elevation-dependant temperatures (Lunt et al. 2016), which can be used to constrain vegetation models (Sepulchre et al. 2006). However, gridded reconstructions of paleo-topographies at regular time intervals have yet to be made publically available. Further, reconstructions of paleo-topography are inherently biased by the fact that sedimentary basins record the subsidence of the Earth′s crust in relatively great detail, while erosion destroys significant information on the temporal dynamics in extent and height of mountain ranges. Biased data collection, partly driven by accessibility issues but largely because sedimentary basins are hosts to the majority of hydrocarbon resources, has further contributed to an imbalance of our understanding of the dynamics of mountain ranges versus sedimentary basins. By now we have a robust understanding of the evolution of oceanic basins (e.g. Müller et al 2016) and continental margins due to decades of data acquisition, research and resource exploration. However, the availability of global and regional crustal-scale data sets (e.g. Pasyanos et al. 2014) on along with improved computational tools allows for potential avenues to construct such paleo-elevation models. These can be further augmented with more detailed regional estimates of paleoelevation in existing databases including thermochronology (e.g. Barns et al. 2012), isotopes studies (e.g. Campani et al. 2012). Effort should be invested in such open access reconstruction to produce a benchmark data set to model terrestrial biodiversity dynamics. Because some fossil data are used to build paleogeographies, those reconstructions are not fully independent from biological data, and this should be considered when interpreting the results. Nevertheless, we anticipate a huge potential of paleogeographies to increase our understanding of spatial biodiversity gradients.

With this paper, we would like to encourage research that bridges paleobiology with macroecology using paleogeographies. Accounting for deep time geological events in spatial models of extant biodiversity is not straightforward and even if our study propose a first illustrative approach, spatial models should be further developed. A new generation of biodiversity models could explicitly model species diversification from the spatio-temporal sequence of environmental conditions (Gotelli et al. 2009). Based on reconstructed paleohabitats, spatial diversification models may track the distribution of lineages in grid cells, as well as their genealogy, which can be compared with multiple empirical evidence, including biodiversity metrics (Gotelli et al. 2009). Leprieur et al. (2016) showed the interest of such an approach to understand tropical marine diversification, which can be compared to current patterns and the fossil records. To this aim, more effort should be invested in producing openly accessible paleoenvironmental maps that can be coupled with mechanistic models to understand the spatial dynamic of species diversification. As stated by Myers and Gillers (1988) “To progress, biogeography must attempt to integrate divergent interests and determine how speciation, adaptation and ecological processes interact with one another and with geology and climate to produce distributional patterns of the world′s biota”. We have now the tools to realize this research agenda and quantify the role of historical habitat dynamics within a fully spatial framework.

## Tables

**Table 1.** Best results of the statistical test of congruence between simulated and observed endemic richness hotspots obtained by taking different values of dispersal distance (d) for terrestrial, freshwater and marine species. The endemic richness was calculated as the sum of the inverse range sizes of all species occurring in a given grid cell. The number of common cells indicate the number of overlapping cells above the 10% threshold (0.1) of the highest value of endemic richness and from the proxy from plate tectonics. The expected hotspots correspond to the mean number of cells of overlap expected from the 999x randomisation.

|  | d | Nb of common cells | Expected hotspots | P value | Threshold | % of common hotspots |
| --- | --- | --- | --- | --- | --- | --- |
| Total Terrestrial | 10 | 435 | 154.47 | 0.001 | 0.1 | 28.79 |
| Amphibians | 30 | 395 | 151.27 | 0.001 | 0.1 | 26.14 |
| Mammals | 30 | 349 | 151.27 | 0.001 | 0.1 | 23.10 |
| Reptiles | 20 | 385 | 151.27 | 0.001 | 0.1 | 25.48 |
| Plants | 30 | 438 | 152.37 | 0.001 | 0.1 | 28.99 |
| Birds | 10 | 449 | 154.47 | 0.001 | 0.1 | 29.72 |
| Total Freshwater | 25 | 407 | 151.27 | 0.001 | 0.1 | 26.94 |
| Crabs | 30 | 440 | 151.57 | 0.001 | 0.1 | 29.12 |
| Mollusks | 20 | 571 | 151.27 | 0.001 | 0.1 | 37.79 |
| Plants | 25 | 606 | 151.27 | 0.001 | 0.1 | 40.11 |
| Fishes | 25 | 335 | 151.27 | 0.001 | 0.1 | 22.17 |
| Total Marine | 10 | 117 | 101.26 | 0.042 | 0.1 | 11.65 |
| Corals | 15 | 132 | 100.56 | 0.001 | 0.1 | 13.15 |
| Conesnails | 15 | 128 | 100.86 | 0.002 | 0.1 | 12.75 |
| Seasnakes | 10 | 55 | 101.46 | 1 | 0.1 | 5.48 |
| Seacucumbers | 25 | 181 | 100.66 | 0.001 | 0.1 | 18.03 |
| Fishes | 10 | 123 | 101.26 | 0.011 | 0.1 | 12.25 |

## Supplementary material

### Supplementary methods

#### Species data and endemic richness

We mapped the global endemic richness for 14 taxonomic group. The metric of endemic richness combines both the level of endemism and species richness as done in Kier et al. (2009). We computed the sum of the inverse range sizes of all species occurring in a given grid cell. The range size of each species was measured as the number of grid cells in which a species occur across the globe. As this method is applicable to gridded distribution data we computed distribution ranges in the form of rasters for 181,603 species on 1° squared grid at global scale. The 14 taxonomic groups can be separated in three categories based on their habitat preferences: (i) for the terrestrial ecosystem we compiled the distribution data for 154,507 species (6,309 amphibians, 5,289 mammals, 128,565 plants, 4,278 reptiles, 10,066 birds); (ii) for freshwater ecosystem we compiled the distribution data for 9,597 species (1,277 crabs, 1,887 mollusks, 135 plants, 6298 fishes); (iii) for marine ecosystem we compiled the distribution data for 17,499 species (844 corals, 632 cone snails, 369 sea cucumbers, 61 sea snakes, 15,593 fishes).

For most of species (amphibians, mammals, reptiles, birds, corals, cone snails, sea cucumbers, crabs, mollusks, freshwater plants, and freshwater fishes) range maps were downloaded from the IUCN website. Species distribution information were compiled in the context of The IUCN Red List assessment (IUCN 2014). The species distribution maps in shapefile format were converted into rasters and aggregated based on a 1° resolution global grid. The distribution information for terrestrial plants was obtained from the Kew worldwide database. We used checklists corresponding to the most detailed “level3” polygons from the Taxonomic Databases Working Group (TDWG; http://www.tdwg.org/, http://www.kew.org/science-conservation/research-data/resources/gis-unit/tdwg-world). Polygons generally correspond to countries, although larger countries are often subdivided into states. For each species, the polygon of the range was rasterized and aggregated on a 1° resolution grid globally.

Fish species occurrence data were obtained from the Ocean Biogeographic Information System (OBIS, http://www.iobis.org). We inventoried 16 238 200 occurrence records from 34.883 entries. We cleaned the data by identifying the synonyms, misspellings and rare species and by restraining it to species present in the marine environment according to FishBase. Synonyms were converted to accepted names. This resulted in a set of 11,503,257 occurrences for 11,345 fish species around the world. We reconstructed distribution maps for each species, defined as the convex polygon surrounding the area where each species was observed. The resulting polygon was divided into four parts across the world to integrate possible discontinuity between the two hemispheres and the Atlantic and Pacific Oceans. We refined the distribution map of each species by removing areas where maximal depths fell outside the minimum or maximum known depth range of the species. Final distribution maps of well-known species were checked visually and reviewed. As the OBIS database did not completely represent the tropical assemblage of fish, we used the GASPAR database at 1° resolution that encompass 6316 coral reef species to complete the database (Pellissier et al. 2014). We crosschecked the databases for the synonymous by using FishBase and we merged them. We obtained a world database containing 15593 fish species that we aggregated on a 1° resolution grid globally.

**Plate tectonic reconstruction**

To investigate whether the dynamic of plate tectonics is associated to the location of hotspots of endemic richness, we generated paleogeographic reconstructions showing the area of exposed land and shallow reefs surface in absolute paleo-positions from the breakup of Gondwana in the Early Cretaceous (140 Ma) to the present in 1 Ma steps. We used the GPlates open source software (http://www.gplates.org;) in conjunction with a newly devised set of digital paleoshoreline positions and a global plate motion model to generate gridded, individual paleogeographic reconstructions for land surface area at any given time step based on geological observations. Those paleogeographies were already used in Leprieur et al. (2016).

#### Modelling of dispersal potential across strait

We developed a model based on land and shallow reef dynamic, which quantifies the amount of potential dispersal into each cell from disconnected patches. The model starts in early Cretaceous times (140 Ma), since layers were not unavailable for deeper time period. The model records at any single point in time and for each cell the number of cells from disconnected patches from which it can receive long distance disperser, which would potentially give rise to a new species. During each million-year time slice, the simulation first define the lands patches separated by at least one cell of sea using an algorithm of cluster splits implemented in the “dbscan” function in the fpc library in R. Then, the model quantifies for each cell, the number of surrounding cells from disconnected land patches below a dispersal distance *d*. A higher number of possible donor cells is assumed to increase the chance of long distance dispersal across a sea barrier. Finally, to accommodate the movement of the continents, the cell values at time t are transferred to the closest cell values at time t+1. Area with high complexity of land patches for extended duration are expected to have the highest potential for allopatric speciation shaping species richness and endemism across barriers. We tested a range of dispersal d (5, 10, 15, 20, 25, 30 [°]).

#### Extinction filter

The previous index highlights the location favourable to species formation through allopatric speciation since the late Cretaceous. However, current species diversity is also the result of extinction, which is supposed to be higher at higher latitude consequently to more pronounced climatic fluctuations. To correct for a gradient in extinction rate associated to latitude and better highlight the location of biodiversity hotspots, we created a filter assuming that survival decreases either linearly or with a Gaussian distribution with latitude. The linear model takes an X-intercept at 0° of latitude and a Y-intercept corresponding to the maximum number in the potential for allopatric speciation. The Gaussian model is centred at 0° of latitude and a range of amplitude parameters were evaluated. The range of values between 15 and 80 were tested for the σ parameter. Finally, we filtered the observed map of potential allopatric speciation to filtering effect of extinction to better highlight the locations of biodiversity hotspots longitudinally.

#### Congruence analysis

We tested whether there is a significant spatial congruence between potential hotspot obtained by the model and the observed endemic richness for the 14 taxonomic groups. We performed a congruence analysis, comparing the observed spatial overlap between hotspots compared to a null-model following Mouillot et al. (2011). For each pairwise comparison, we calculated the number of cells recorded as a hotspot (10% of highest values) for both observed and modelled and the degree of independence between the two hotspot ensembles. To assess the significance of observed overlaps from that obtained by chance, we computed a randomization procedure. We randomly permuted (x 999) values contained in cells of one of the two considered variables. The percentage of cell overlaps was estimated for each permutation. In this analysis, the number of common hotspots represents the number of cells that are recorded as a hotspot for both observed and predicted patterns. To avoid the overestimation of hotspot in arid areas, we excluded the cells classified as desert from this analysis. The expected hotspots corresponding to the independence between the two hotspot ensembles.

## Supplementary table

**Table S1:** Best results of the test of congruence between simulated speciation hotspots and observed endemic richness hotspots obtained by taking different values of dispersal distance (d) for terrestrial, freshwater and marine species while also considering a decreasing linear degree of extinction from the pole to the equator. The endemic richness was calculated as the sum of the inverse range sizes of all species occurring in a given grid cells. The number of common cells indicate the number of overlapping cells above the 10% threshold (0.1) of the highest value of endemic richness and from the proxy from plate tectonics. The expected hotspots correspond to the mean number of cells of overlap expected from the 999x randomisation. The spearman correlations and associated p-values are also provided.

|  | d | Nb of<br>common<br>cells | Expected<br>hotspots | P<br>value | Threshold | % of<br>common<br>hotspots | ρ de<br>spearman | P value |
| --- | --- | --- | --- | --- | --- | --- | --- | --- |
| Total terrestrial | 15 | 613 | 151.27 | 0.001 | 0.1 | 40.57 | 0.36 | < 0.001 |
| Amphibians | 15 | 517 | 151.27 | 0.001 | 0.1 | 34.22 | 0.36 | < 0.001 |
| Mammals | 15 | 513 | 151.27 | 0.001 | 0.1 | 33.95 | 0.38 | < 0.001 |
| Reptiles | 15 | 487 | 151.27 | 0.001 | 0.1 | 32.23 | 0.35 | < 0.001 |
| Plants | 15 | 628 | 152.37 | 0.001 | 0.1 | 41.56 | 0.34 | < 0.001 |
| Birds | 15 | 633 | 151.27 | 0.001 | 0.1 | 41.89 | 0.39 | < 0.001 |
| Total freshwater | 25 | 359 | 151.27 | 0.001 | 0.1 | 23.76 | 0.38 | < 0.001 |
| Crabs | 25 | 631 | 151.57 | 0.001 | 0.1 | 41.76 | 0.51 | < 0.001 |
| Molluscs | 30 | 395 | 151.27 | 0.001 | 0.1 | 26.14 | 0.29 | < 0.001 |
| Plants | 30 | 532 | 151.27 | 0.001 | 0.1 | 35.21 | 0.35 | < 0.001 |
| Fishes | 25 | 354 | 151.27 | 0.001 | 0.1 | 23.43 | 0.21 | < 0.001 |
| Total marine | 30 | 219 | 100.56 | 0.001 | 0.1 | 21.81 | 0.56 | < 0.001 |
| Corals | 30 | 423 | 100.56 | 0.001 | 0.1 | 42.13 | 0.62 | < 0.001 |
| Conesnails | 30 | 285 | 100.86 | 0.001 | 0.1 | 28.39 | 0.62 | < 0.001 |
| Seasnakes | 25 | 277 | 100.76 | 0.001 | 0.1 | 27.59 | 0.46 | < 0.001 |
| Seacucumbers | 25 | 315 | 100.66 | 0.001 | 0.1 | 31.37 | 0.49 | < 0.001 |
| Fishes | 30 | 212 | 100.56 | 0.001 | 0.1 | 21.12 | 0.44 | < 0.001 |

**Table S2:** Best results of the test of congruence between simulated and observed endemic richness hotspots obtained by taking different values of dispersal distance (d) for terrestrial, freshwater and marine species while considering a decreasing Gaussian function of extinction from the pole to the equator. The ndemic richness was calculated as the sum of the inverse range sizes of all species occurring in a given grid cells. The parameter μ of the Gaussian function was set to 0 to represent a decreasing extinction from the pole to the equator. All values between 15 and 80 were tested for the σ parameter. The number of common cells indicate the number of overlapping cells above the 10% threshold (0.1) of the highest value of endemic richness and from the proxy from plate tectonics. The expected hotspots correspond to the mean number of cells of overlap expected from the 999x randomisation. The spearman correlations and associated p-values are also provided.

| | d | $\sigma$ | Nb of common cells | Expected hotspot | P value | Threshold | % of common hotspots | $\rho$ de spearman | P value |
| --- | --- | --- | --- | --- | --- | --- | --- | --- | --- |
| Total Terrestrial | 5 | 45 | 713 | 151.47 | 0.001 | 0.1 | 47.19 | 0.37 | < 0.001 |
| Amphibian | 5 | 45 | 592 | 151.47 | 0.001 | 0.1 | 39.18 | 0.37 | < 0.001 |
| Mammals | 20 | 18 | 611 | 151.27 | 0.001 | 0.1 | 40.44 | 0.46 | < 0.001 |
| Reptiles | 5 | 57 | 639 | 151.37 | 0.001 | 0.1 | 42.29 | 0.43 | < 0.001 |
| Plants | 5 | 45 | 702 | 152.57 | 0.001 | 0.1 | 46.46 | 0.39 | < 0.001 |
| Birds | 5 | 45 | 724 | 151.47 | 0.001 | 0.1 | 47.92 | 0.41 | < 0.01 |
| Total Freshwater | 25 | 79 | 384 | 151.27 | 0.001 | 0.1 | 25.41 | 0.68 | < 0.01 |
| Crabs | 5 | 45 | 738 | 151.77 | 0.001 | 0.1 | 48.84 | 0.42 | < 0.001 |
| Molluscs | 25 | 79 | 498 | 151.27 | 0.001 | 0.1 | 32.96 | 0.35 | < 0.001 |
| Plants | 25 | 79 | 595 | 151.27 | 0.001 | 0.1 | 39.38 | 0.44 | < 0.001 |
| Fishes | 30 | 20 | 415 | 151.27 | 0.001 | 0.1 | 27.46 | 0.17 | < 0.001 |
| Total Marine | 30 | 20 | 248 | 100.56 | 0.001 | 0.1 | 24.70 | 0.36 | < 0.001 |
| Corals | 30 | 20 | 532 | 100.56 | 0.001 | 0.1 | 52.99 | 0.61 | < 0.001 |
| Conesnails | 30 | 32 | 308 | 100.86 | 0.001 | 0.1 | 30.68 | 0.66 | < 0.001 |
| Seasnakes | 15 | 17 | 387 | 100.86 | 0.001 | 0.1 | 38.55 | 0.47 | < 0.001 |
| Seacucumbers | 30 | 32 | 329 | 100.66 | 0.001 | 0.1 | 32.77 | 0.5 | < 0.001 |
| Fishes | 30 | 20 | 237 | 100.56 | 0.001 | 0.1 | 23.61 | 0.37 | < 0.001 |

## Supplementary figures

**Figure S1:**
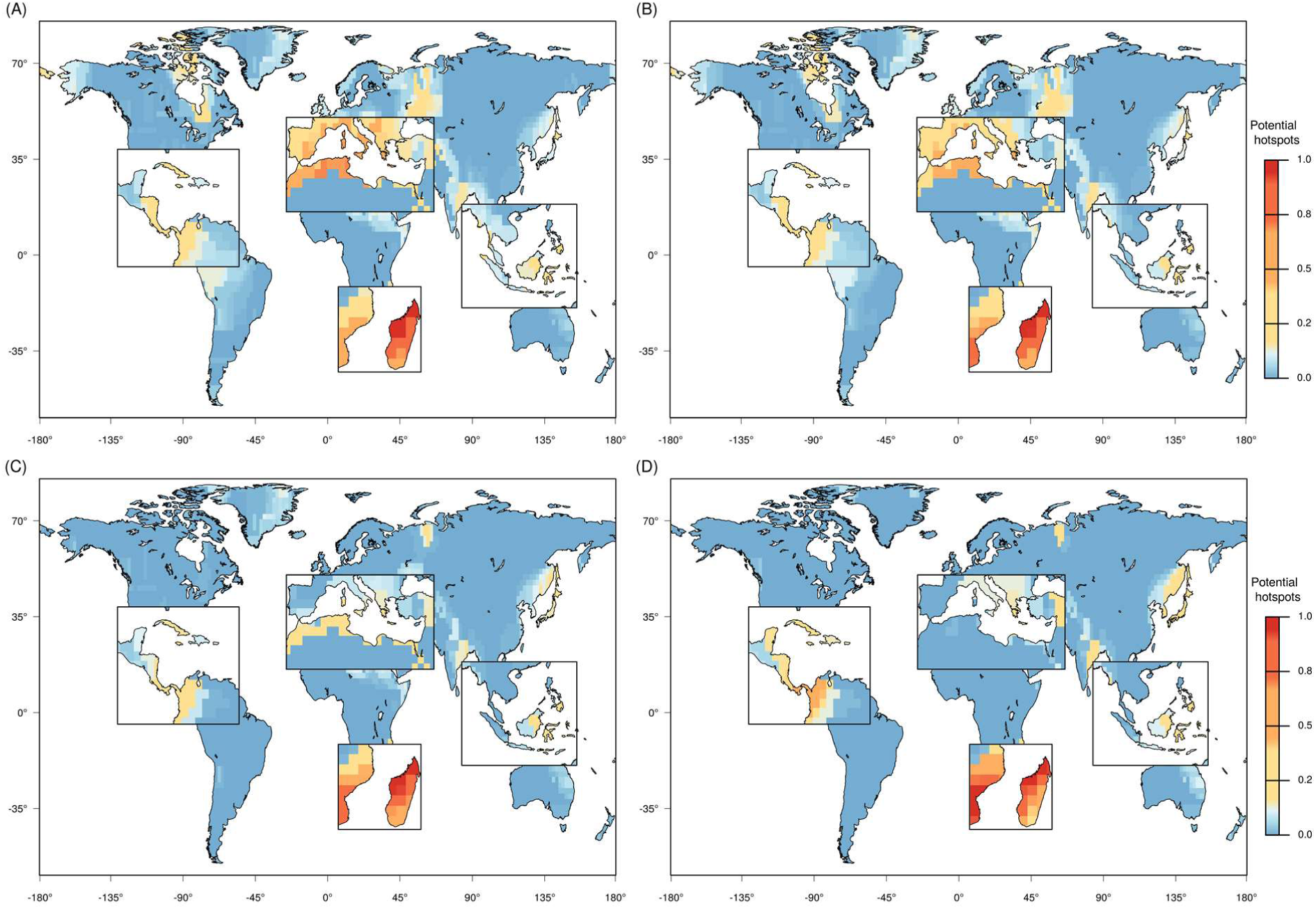
Outputs of the simulations with different starting times (A) 100Mya, (B), 80Mya, (C) 60Mya, (D) 40Mya. The map of the final step of the simulation starting in the Cretaceous (140Mya) was highly correlated to the outputs using more recent starting dates (e.g. for *d*=10, Pearson′s correlations with 100Mya: cor=0.92, 80 Mya: cor=0.88, 60 Mya cor=0.72, 40 Mya cor=0.66). Specifically, the spatial distribution of high values is conserved, only the strength of the effect in Madagascar relative to the other hotspots tend to increase with a more recent start.

**Figure S2:**
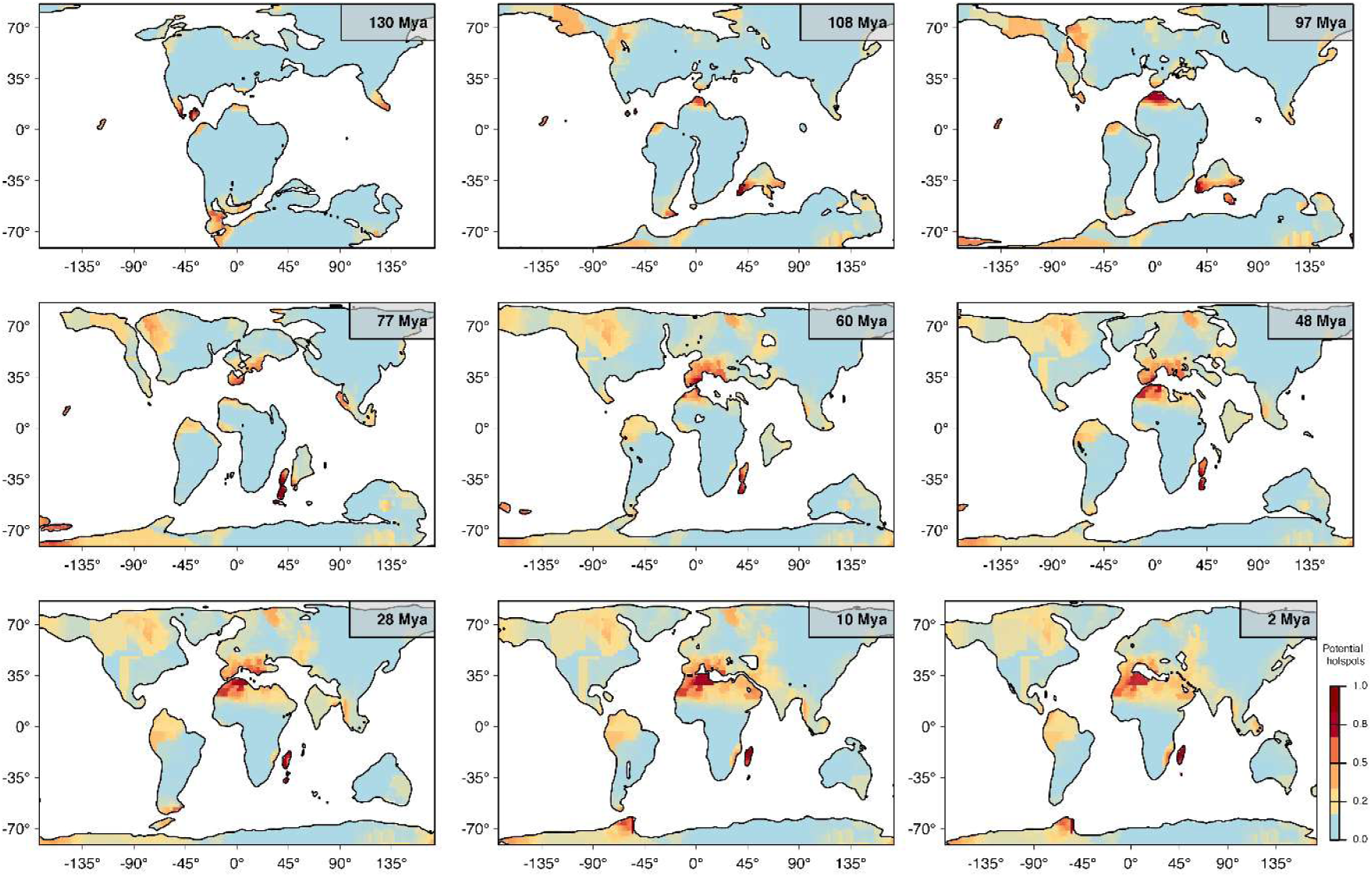
Evolution through time of the favourable configurations of paleogeographies for speciation through allopatric speciation across sea straits, here shown with the parameter value of dispersal of *d*=10 for 9 time periods across the past 130 Mya.

**Figure S3:**
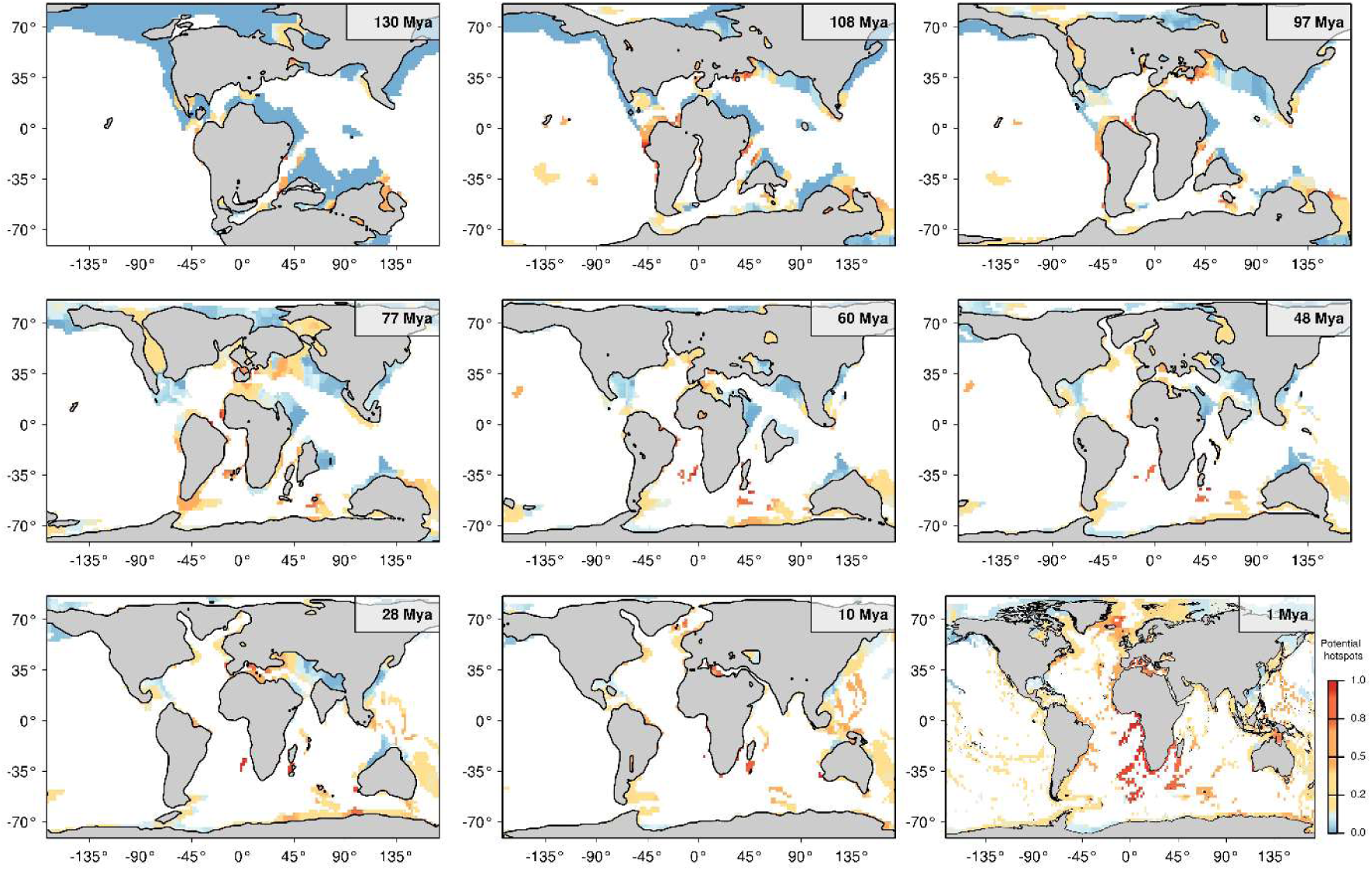
Evolution through time of the favourable configurations of marine shallow marine reef habitats for speciation through allopatric speciation across deep sea, here shown with a dispersal *d*=10 for 9 time periods across the past 130 Mya. The region south west of Africa has particularly high simulated values since from the separation of South America and Africa, it was characterized by fragmented reef patches, which fostered an increase in simulated values through time.

**Figure S4:**
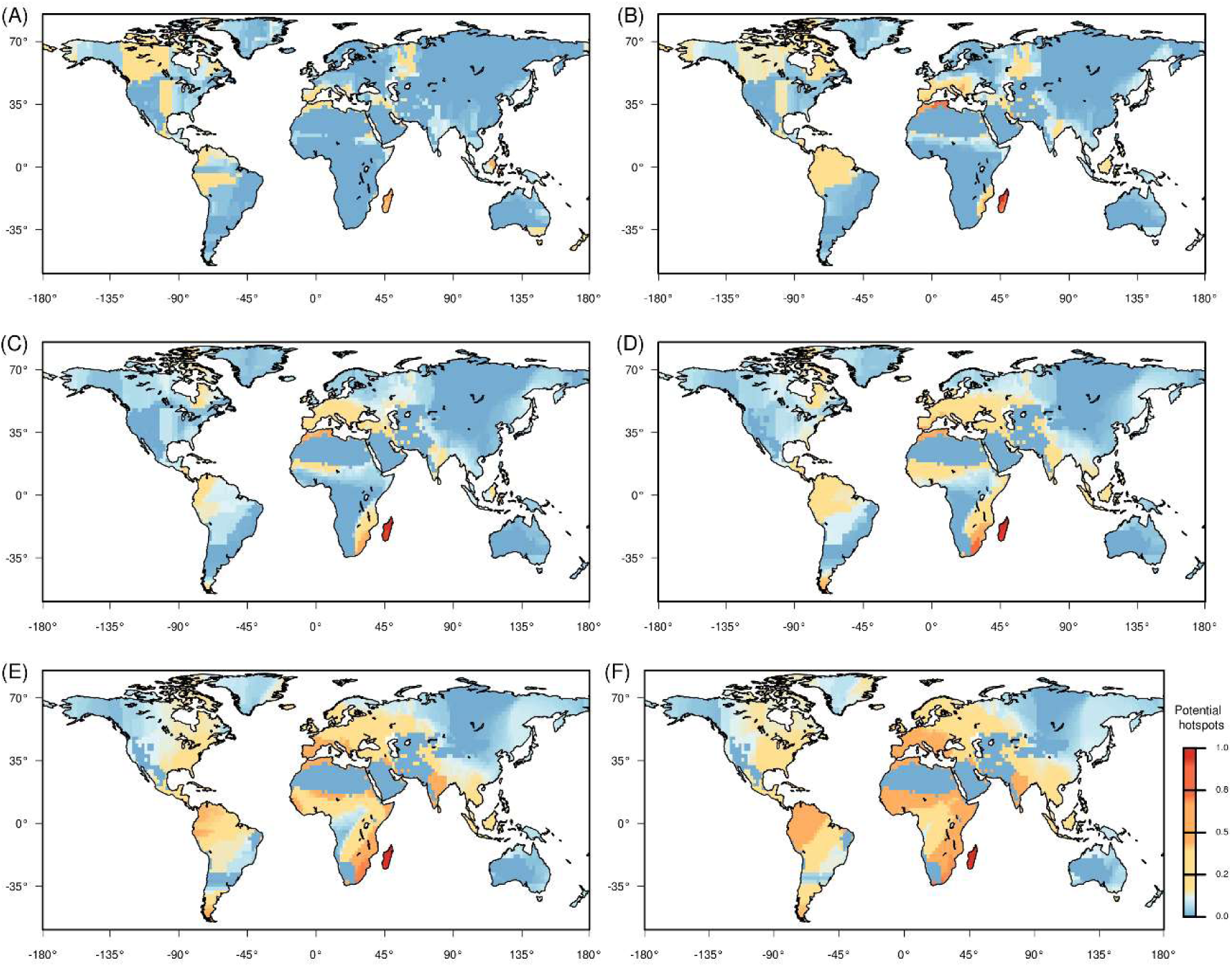
Maps of the terrestrial areas predicted by the model under allopatric speciation hypothesis and dispersal across sea strait considering several values for the dispersal distance parameter: (A) d = 5; (B) d = 10; (C) d = 15; (D) d = 20; (E) d = 25; (F) d = 30.

**Figure S5:**
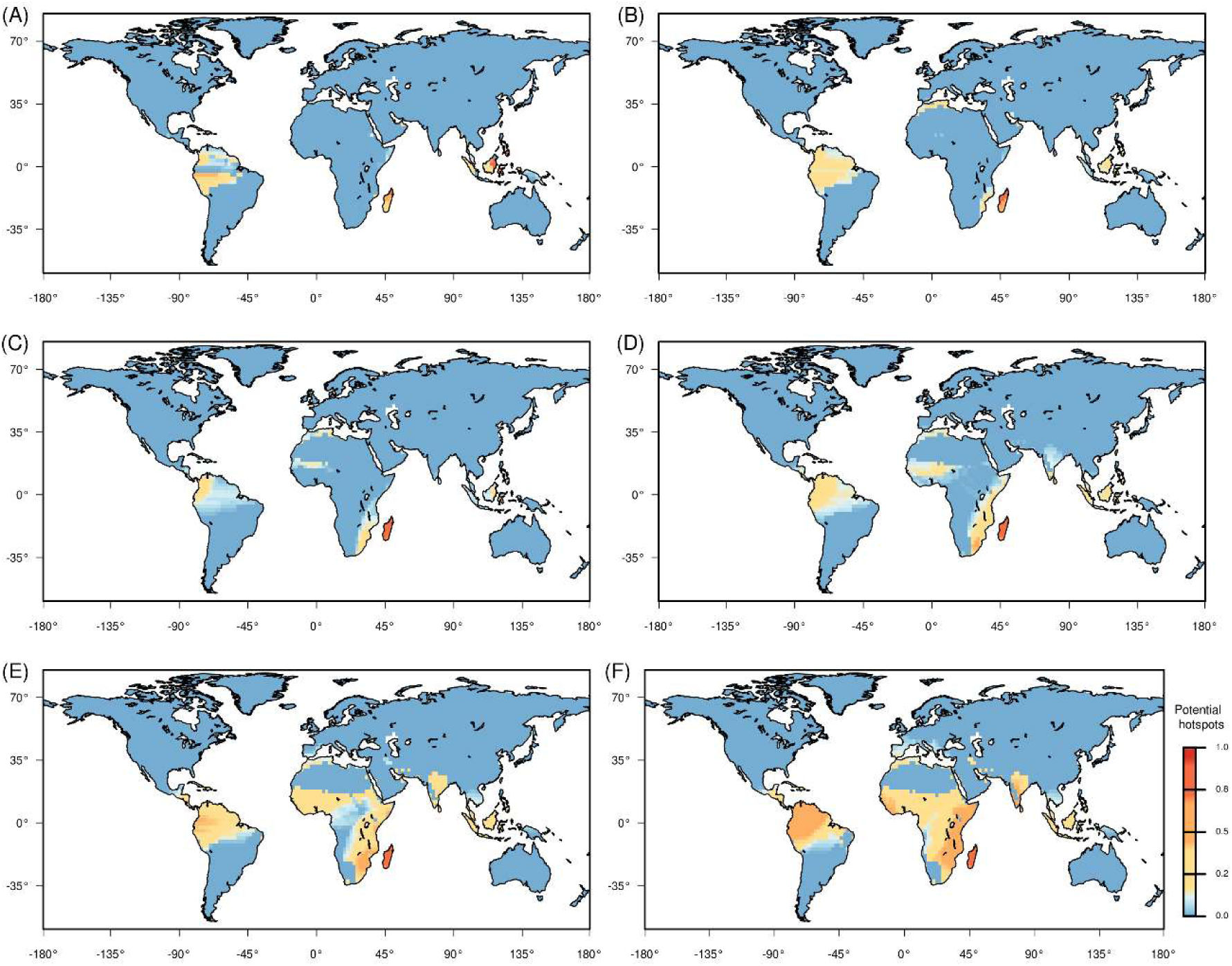
Maps of the terrestrial areas favourable for allopatric speciation predicted by the model for several values of dispersal distance parameter and with considering a decreasing linear degree of extinction from the pole to the equator: (A) d = 5; (B) d = 10; (C) d = 15; (D) d = 20; (E) d = 25; (F) d = 30.

**Figure S6:**
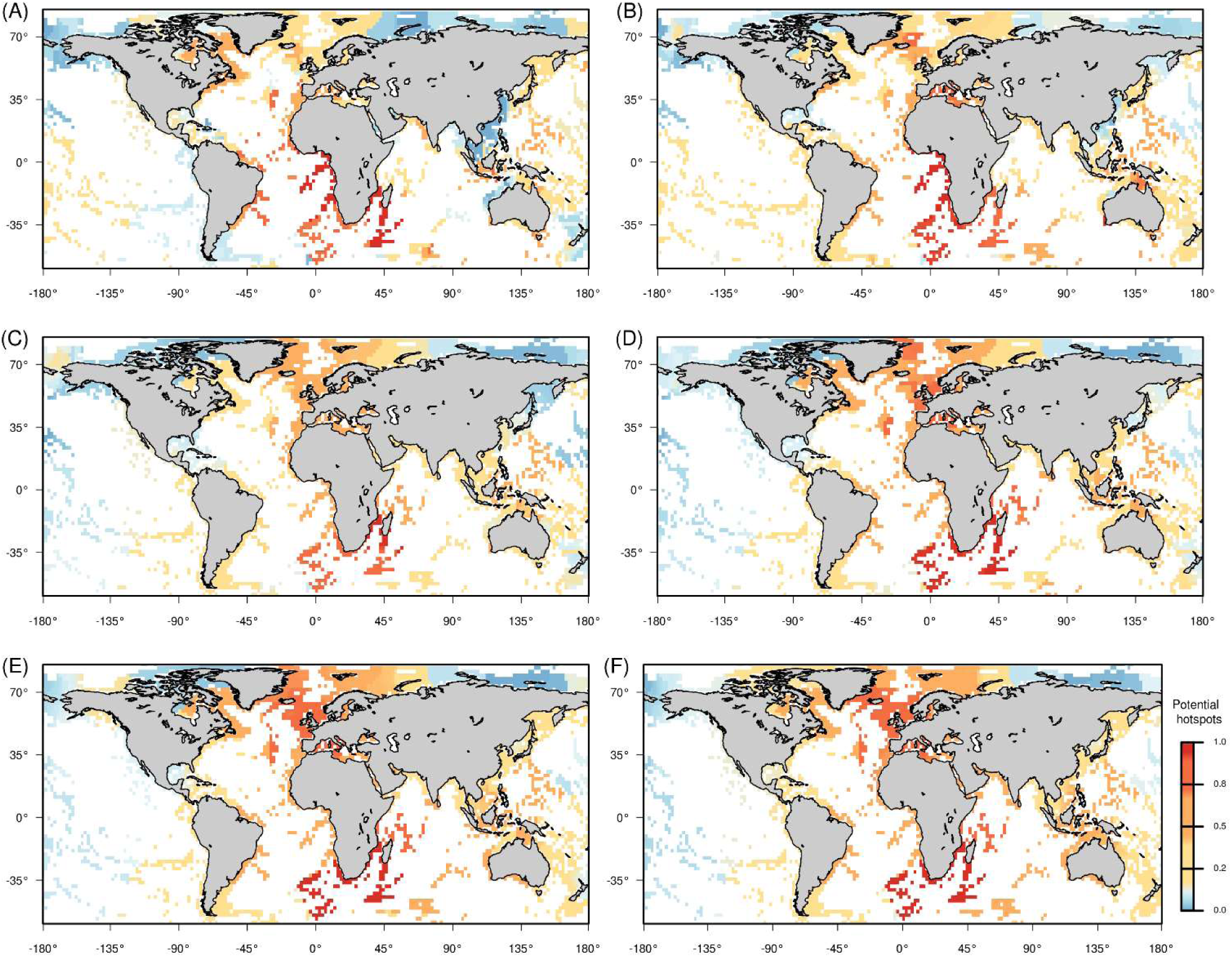
Maps of the coastal marine areas favourable for allopatric speciation predicted by the model for several values of dispersal distance parameter with considering non extinction from the pole to the equator: (A) d = 5; (B) d = 10; (C) d = 15; (D) d = 20; (E) d = 25; (F) d = 30.

**Figure S7:**
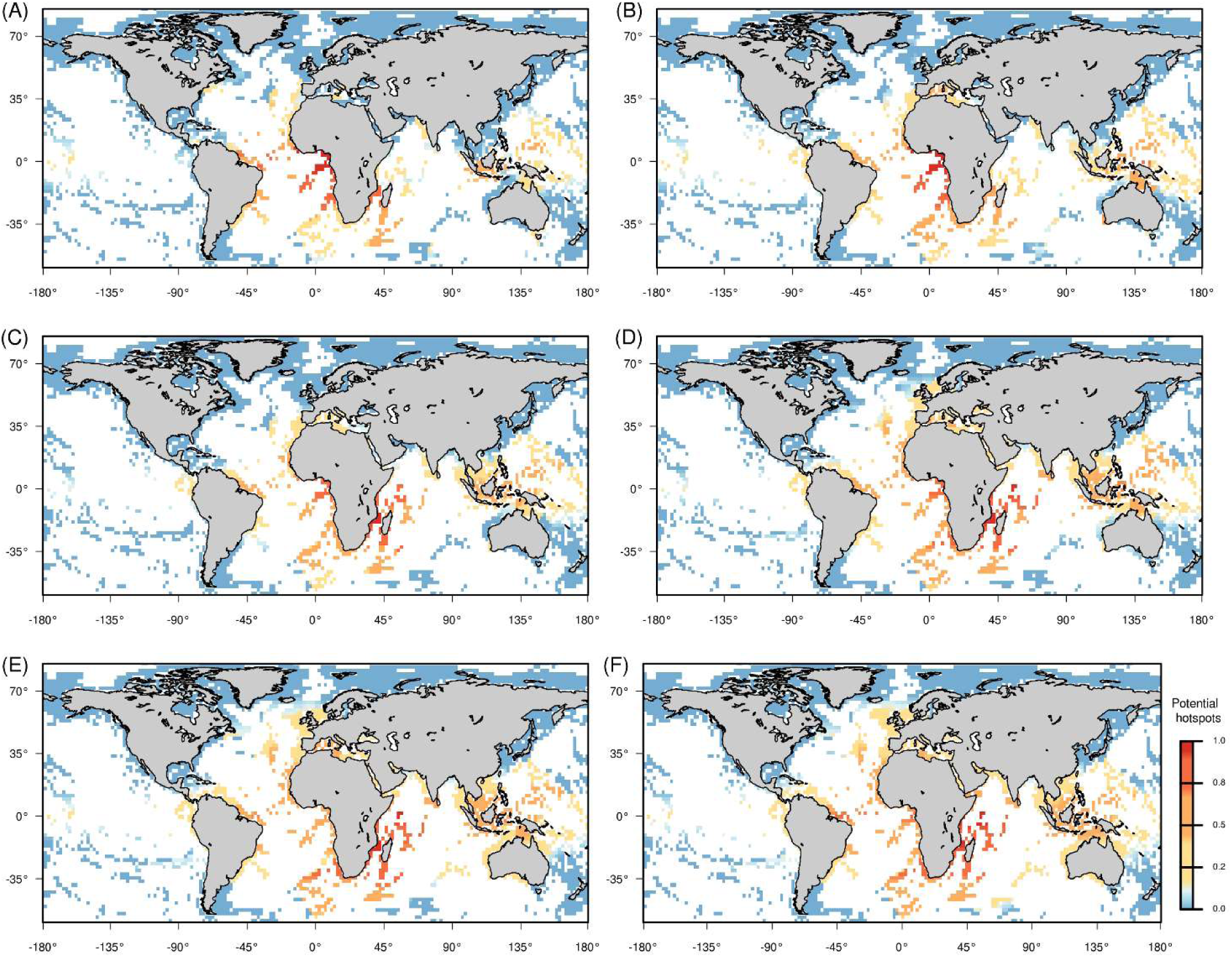
Maps of the coastal marine areas favourable for allopatric speciation predicted by the model for several values of dispersal distance parameter and with considering a decreasing linear degree of extinction from the pole to the equator: (A) d = 5; (B) d = 10; (C) d = 15; (D) d = 20; (E) d = 25; (F) d = 30.

**Figure S8:**
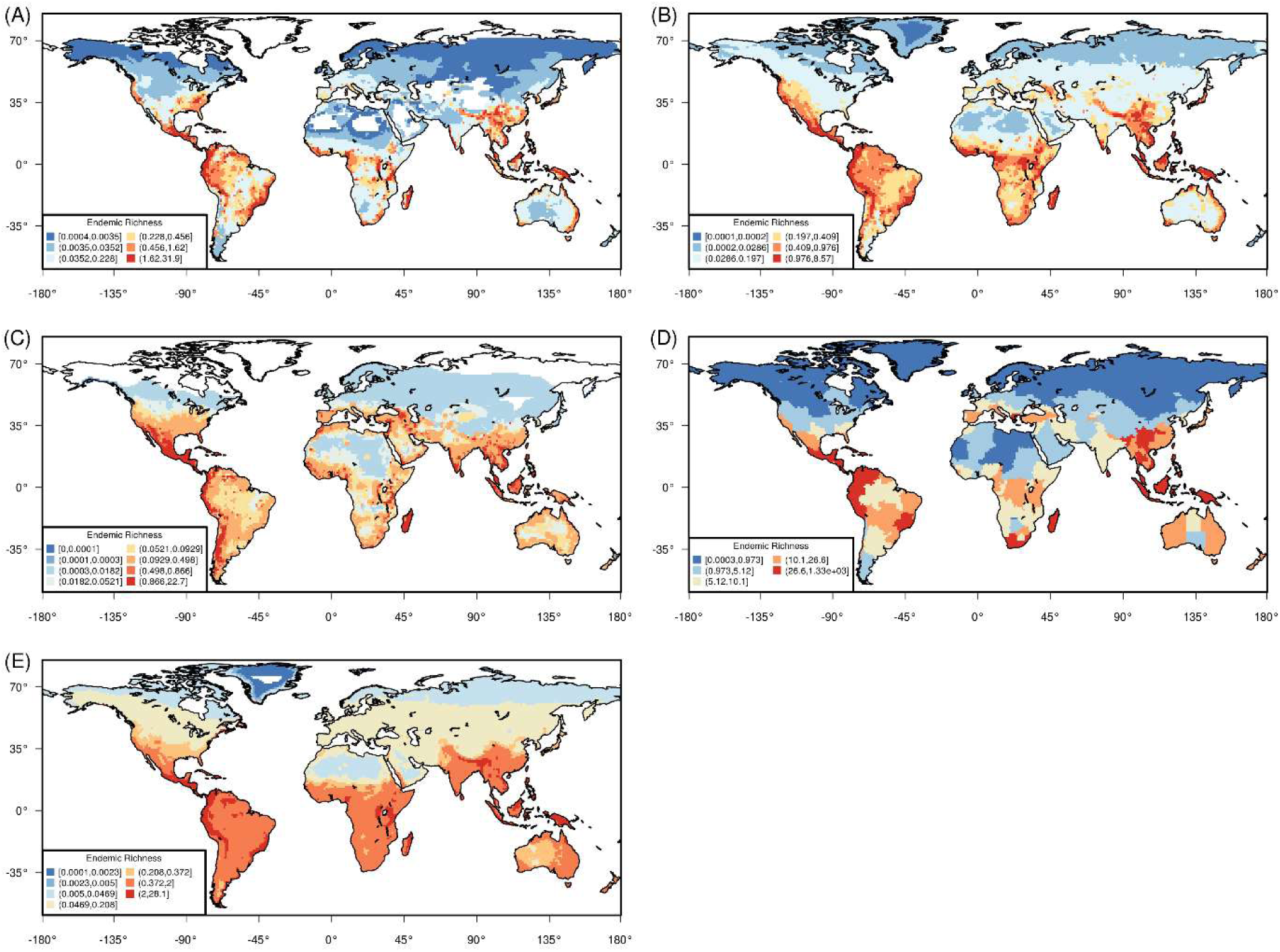
Map of the terrestrial endemic richness calculated as the sum of the inverse range size of all species occurring in a given cells for terrestrial groups: (A) Amphibian n = 6309, (B) Mammals n = 5289, (C) Reptiles n = 4278, (D) Plants n = 128565, (E) Birds n= 10066.

**Figure S9:**
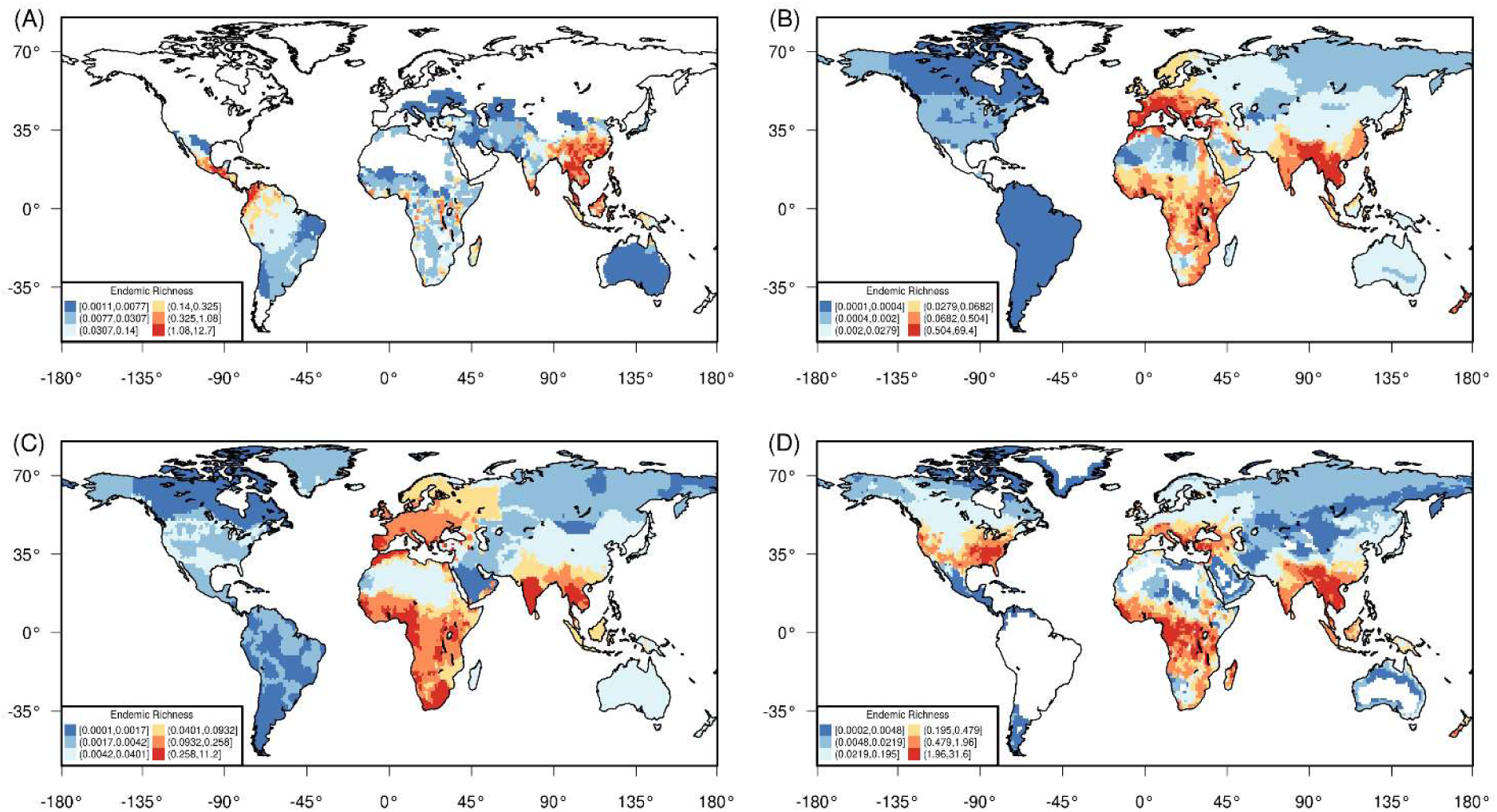
Map of the freshwater endemic richness calculated as the sum of the inverse range size of all species occurring in a given cells for terrestrial freshwater groups (A) Crabs n = 1277, (B) Mollusc n=1887, (C) Plants n= 135 and (D) Fishes n = 6298. Data for freshwater species are not complete in some regions of the world, especially mollusc in South Americas.

**Figure S10:**
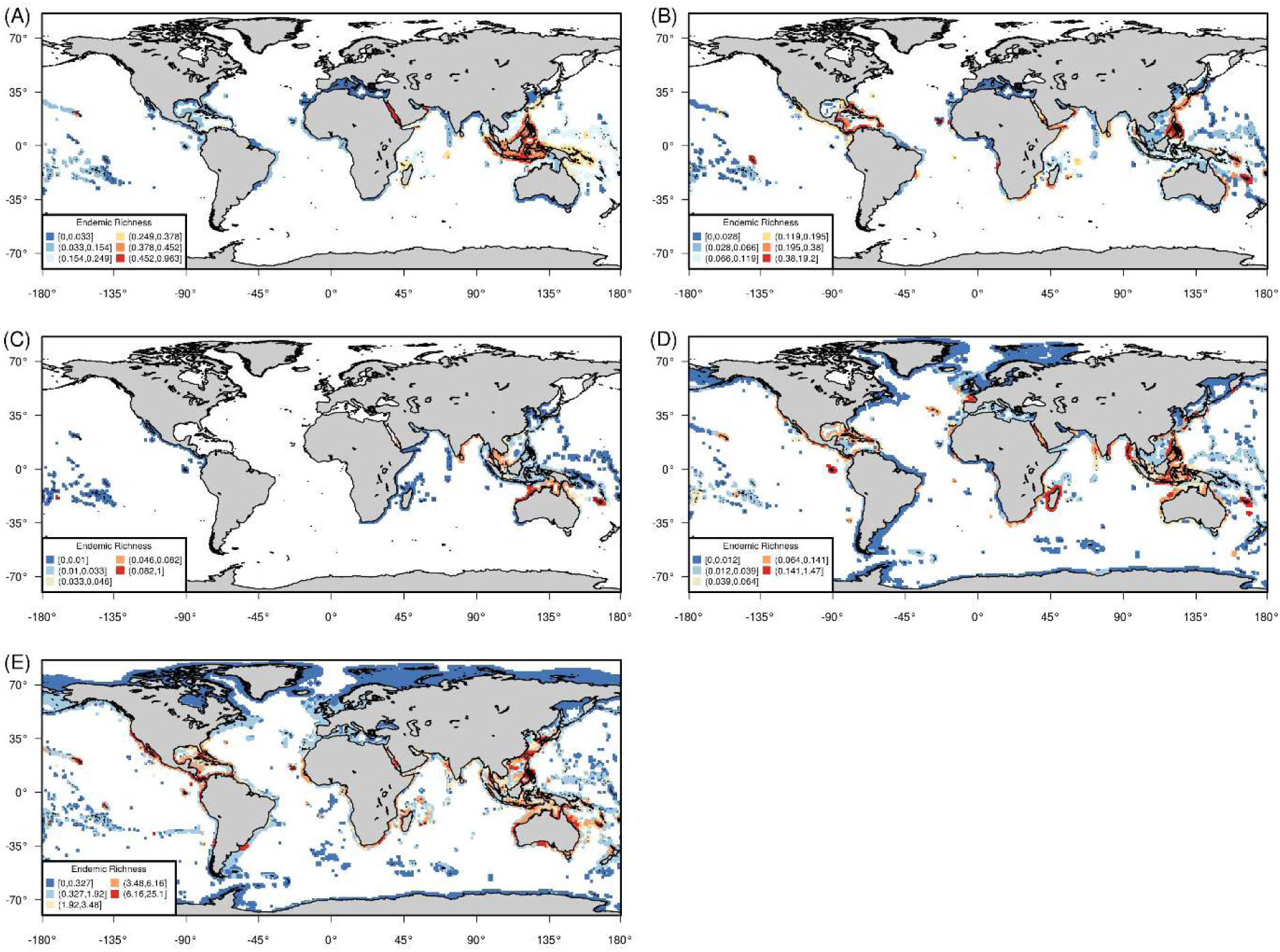
Map of the marine endemic richness calculated as the sum of the inverse range size of all species occurring in a given cells for marine species groups (A) Corals n = 844, (B) Conesnails n = 632, (C) Seasnakes n = 61, (D) Seacucumbers n = 369 and (E) Fishes n = 15593.

**Figure S11:**
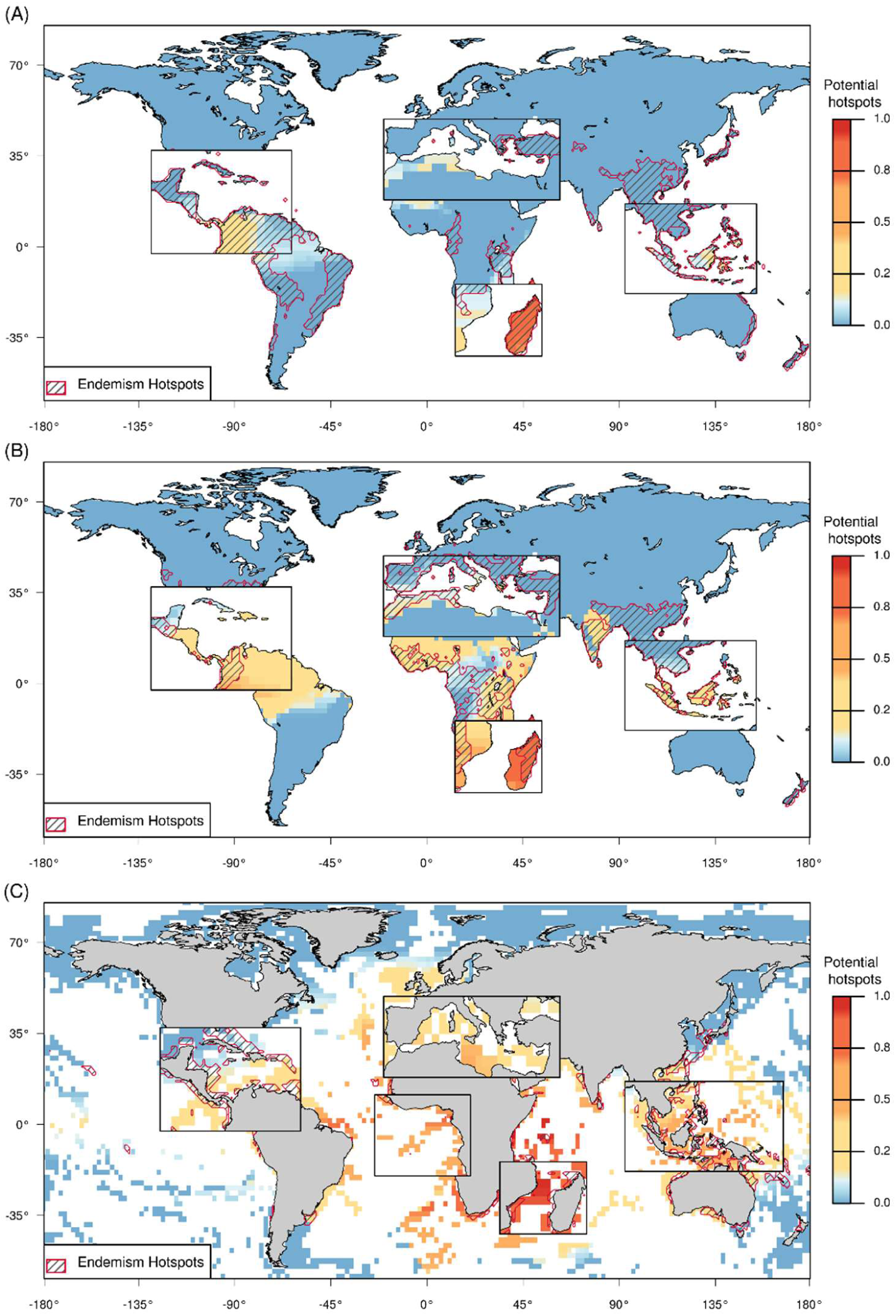
Map of the potential hotspots for allopatric speciation across sea strait for the terrestrial (A) and freshwater species (B) and across reef separated by deep water for marine species (C) predicted by the model with d = 15 for A, d = 25 for B and d = 30 for C with considering decreasing linear extinction from the pole to the equator. Relevant areas are highlighted by zoom plot and compared to the hotspots (10% of the highest values) of endemic richness (red contour shape and black hatches).

**Figure S12:**
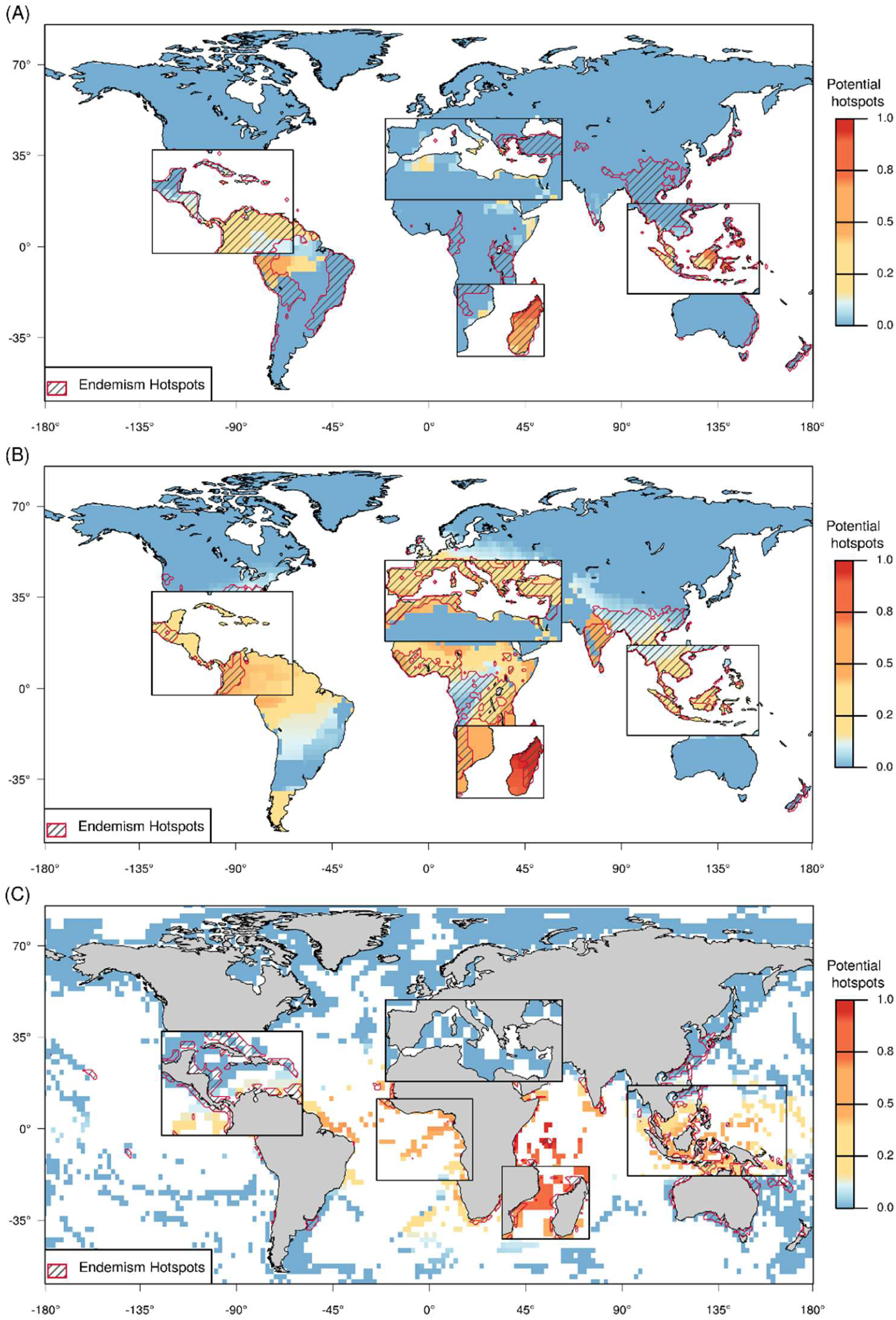
Map of the potential hotspots for allopatric speciation across sea strait for the terrestrial (A) and freshwater species (B) and across shallow reefs separated by deep water for marine species (C) predicted by the model with d = 5 and *σ* = 45 for A, d = 25 *σ* = 79 for B and d = 30 *σ* = 20 for C with considering decreasing a Gaussian extinction function from the pole to the equator (*μ* was set at 0 for all simulations). Relevant areas are highlighted by zoom plot and compared to the hotspots (10% of the highest values) of endemic richness (red contour shape and black hatches).

**Figure S13:**
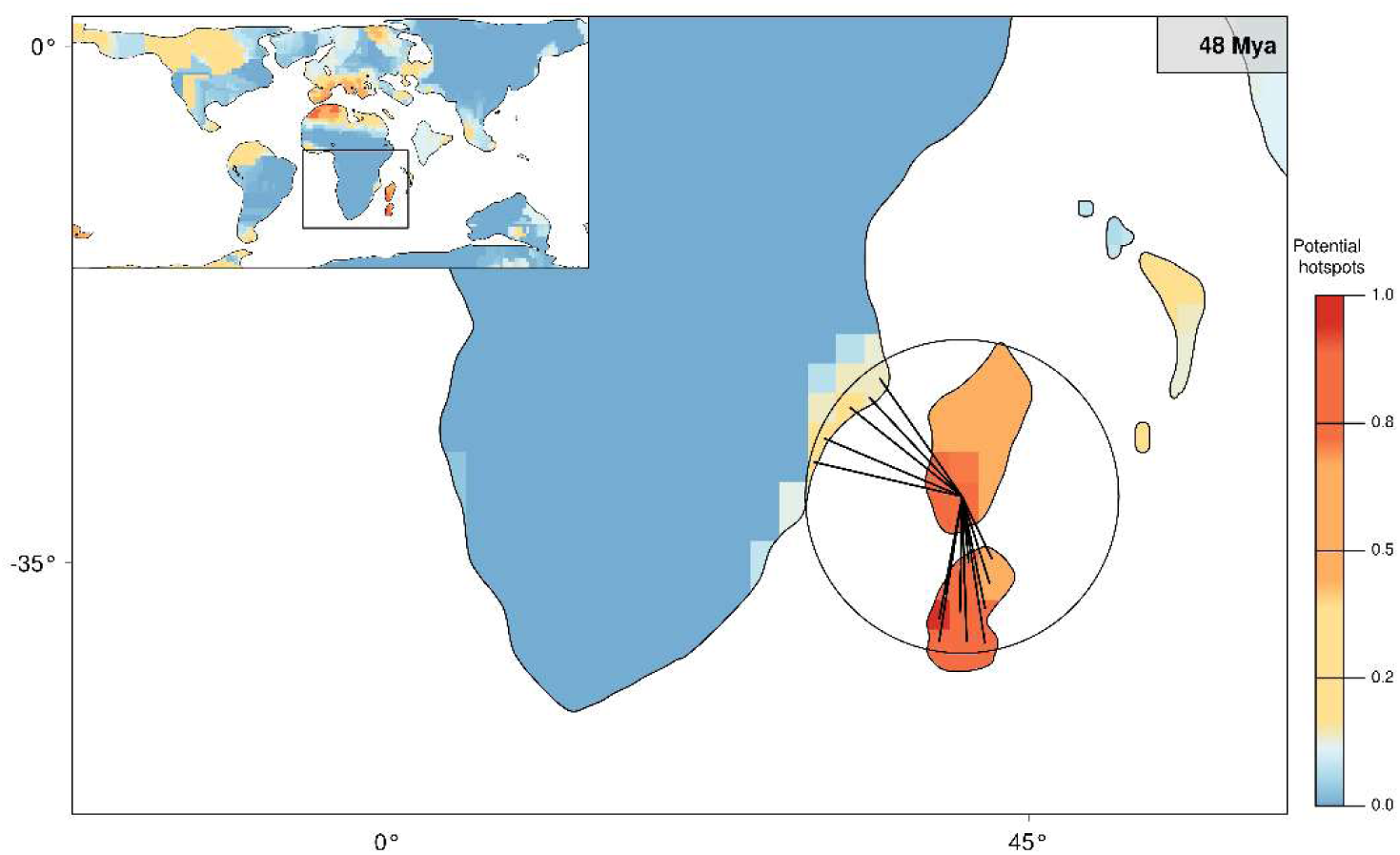
Illustration of the functioning of the spatial model highlighting the position of potential hotspots for allopatric speciation through time. For any given cell, the model computes and cumulates the number of donor cells, which are at a distance >d. In this example, the number of donor cells is computed for a receiver cell in Madagascar. Those values are then cumulated through time.

## References

Ali, J. R. and Huber, M. 2010. Mammalian biodiversity on Madagascar controlled by ocean currents. – Nature 463: 653–656.

Ali, J. R. and Aitchison, J. C. 2014. Exploring the combined role of eustasy and oceanic island thermal subsidence in shaping biodiversity on the Galapagos. – J. Biogeogr. 41: 1227–1241.

Ali, J. R. and Aitchison, J. C. 2008. Gondwana to Asia: plate tectonics, paleogeography and the biological connectivity of the Indian sub-continent from the Middle Jurassic through latest Eocene (166–35 Ma). – Earth-Science Reviews, 88: 145–166.

Bacon, C. D. et al. 2013. Geographic and taxonomic disparities in species diversity: Dispersal and diversification rates across Wallace’s Line. – Evolution 67: 2058–2071.

Bagley, J. C. and Johnson, J. B. 2014. Phylogeography and biogeography of the lower Central American Neotropics: diversification between two continents and between two seas. – Biol. Rev. 89: 767–790.

Badgley, C. 2010. Tectonics, topography, and mammalian diversity. – Ecography, 33: 220–231.

Barnes, J. B. et al. 2012. Linking orography, climate, and exhumation across the central Andes. – Geology, 40:1135–1138.

Bellwood, D. R. et al. 2005. Environmental and geometric constraints on Indo-Pacific coral reef biodiversity. – Ecol. Lett., 8: 643–651.

Bidegaray-Batista, L., & Arnedo, M. A. 2011. Gone with the plate: the opening of the Western Mediterranean basin drove the diversification of ground-dweller spiders. – BMC Evol. Biol., 11, 1.

Briand, F., and Cohen, J. E. 1987. Environmental correlates of food chain length. – Science, 238, 956–960.

Briggs, J. C. 2003. The biogeographic and tectonic history of India. – J. Biogeogr., 30, 381–388.

Campani, M. et al. 2012. Miocene paleotopography of the Central Alps. – Earth Planet. Sci. Lett, 337, 174–185.

Cracraft, J. 1973. Continental drift, paleoclimatology, and the evolution and biogeography of birds. – J. Zool. 169: 455–543.

Condamine, F. L. et al. 2012. What causes latitudinal gradients in species diversity? Evolutionary processes and ecological constraints on swallowtail biodiversity. – Ecol. Lett. 15, 267–277.

Couvreur, T. L. P. 2015. Odd man out: why are there fewer plant species in African rain forests? – Plant Syst. Evol. 301: 1299–1313.

Cowie, R. H. and Holland, B. S. 2006. Dispersal is fundamental to biogeography and the evolution of biodiversity on oceanic islands. – J. Biogeogr. 33: 193–198.

Currie, D. J. 1991. Energy and large-scale patterns of animal- and plant-species richness. – Am. Nat. 137: 27–49.

Davies, R. G. et al. 2007. Topography, energy and the global distribution of bird species richness. – Proc. R. Soc. Biol. Sci. 274: 1189–1197.

Dynesius, M., and Jansson, R. 2000. Evolutionary consequences of changes in species’ geographical distributions driven by Milankovitch climate oscillations. – Proc. Natl Acad. Sci. USA 97: 9115–9120.

Flessa, K. W. (1980). Biological effects of plate tectonics and continental drift. BioScience, 30: 518– 523.

Futuyma, D. J., & Mayer, G. C. 1980. Non-allopatric speciation in animals. – System. Biol. 29: 254–271.

Gotelli, N. J. et al. 2009. Patterns and causes of species richness: a general simulation model for macroecology. – Ecol. Lett. 12, 873–886.

Graham, C. H. et al. 2006. Habitat history improves prediction of biodiversity in rainforest fauna. – Proc. Natl Acad. Sci. USA 103: 632–636.

Gillespie, R. G. and Roderick, G. K. 2014. Evolution: Geology and climate drive diversification. – Nature 509: 297–298.

Guzmán, B. and Vargas, P. 2009. Long-distance colonization of the Western Mediterranean by Cistus ladanifer (Cistaceae) despite the absence of special dispersal mechanisms. – J. Biogeogr. 36: 954–968.

Hawkins, B. A. et al. 2003. Energy, water, and broad-scale geographic patterns of species richness. – Ecology, 84, 3105–3117.

Hedges, S. B. et al. 1996. Continental breakup and the ordinal diversification of birds and mammals. – Nature 381: 226–229.

Heine, C. et al. 2015. Evaluating global paleoshoreline models for the Cretaceous and Cenozoic. – Aust. J. Earth Sci. 62: 275–287.

Hoorn, C. et al. 2010. Amazonia through time: Andean uplift, climate change, landscape evolution, and biodiversity. – Science 330: 927–31.

Hodge, J. R., and Bellwood, D. R. 2016. The geography of speciation in coral reef fishes: the relative importance of biogeographical barriers in separating sister-species. – J. Biogeogr. 43:1324–1335

IUCN 2014. The IUCN Red List of Threatened Species. Version 2014.1. http://www.iucnredlist.org. Downloaded on 03/2016.

Keith, S. A. et al. 2013. Faunal breaks and species composition of Indo-Pacific corals: the role of plate tectonics, environment and habitat distribution. – Proc. R. Soc. Biol. Sci. 280: 20130818.

Kreft, H. and Jetz, W. 2007. Global patterns and determinants of vascular plant diversity. – Proc. Natl Acad. Sci. USA 104: 5925–5930.

Jetz, W. and Fine, P. V. 2012. Global gradients in vertebrate diversity predicted by historical area-productivity dynamics and contemporary environment. – PLoS Biol. 10: e1001292.

Jetz, W. et al. 2012. The global diversity of birds in space and time. – Nature 491: 444–448.

Kay, R. F. et al. 1997. Primate species richness is determined by plant productivity: implications for conservation. – Proc. Natl Acad. Sci. USA 94: 13023–13027.

Kerkhoff, A. J. et al. 2014. The latitudinal species richness gradient in New World woody angiosperms is consistent with the tropical conservatism hypothesis. – Proc. Natl Acad. Sci. USA 111: 8125–8130.

Kier, G. et al. 2009. A global assessment of endemism and species richness across island and mainland regions. – Proc. Natl. Acad. Sci. U. S. A. 106: 9322–9327.

Latham, R. E. and Ricklefs, R. E. 1993. Global patterns of tree species richness in moist forests: energy-diversity theory does not account for variation in species richness. – Oikos 67: 325–333.

Lavergne, S. et al. 2013. In and out of Africa: how did the Strait of Gibraltar affect plant species migration and local diversification? – J. Biogeogr. 40: 24–36.

Leprieur, F. et al. 2016. Plate tectonics drive tropical reef biodiversity dynamics. – Nature Com. 7.

Li, J. T. et al. 2013. Diversification of rhacophorid frogs provides evidence for accelerated faunal exchange between India and Eurasia during the Oligocene. – Proc. Natl Acad. Sci. USA 110: 3441–3446.

Lunt, D. J. et al. 2016. Palaeogeographic controls on climate and proxy interpretation. – Climate of the Past 12: 1181–1198.

Magri, D. et al. 2007. The distribution of Quercus suber chloroplast haplotypes matches the palaeogeographical history of the western Mediterranean. – Molecular Ecology 16: 5259– 5266.

Meredith, R. W. et al. 2011. Impacts of the cretaceous terrestrial revolution and KPg extinction on mammal diversification. – Science, 334: 521–524.

Mittelbach, G. G. et al. 2007. Evolution and the latitudinal diversity gradient: speciation, extinction and biogeography. – Ecol. Lett. 10: 315–331.

Mouillot, D. et al. 2011. Protected and threatened components of fish biodiversity in the Mediterranean Sea. – Curr. Biol. 21: 1044–1050.

Müller, R. D. 2008. Long-term sea-level fluctuations driven by ocean basin dynamics. – Science 1357: 1357–1363.

Müller, R.D. et al. Ocean basin evolution and global-scale plate reorganization events since Pangea breakup. – Ann. Rev. Earth Plan. Sci. 44: 107–138

Myers, A. A., and Giller, P. S. 1988. Process, pattern and scale in biogeography. In Analytical Biogeography (pp. 3–12). Springer Netherlands.

Near, T. J. et al. 2012. Ancient climate change, antifreeze, and the evolutionary diversification of Antarctic fishes. – Proc. Natl Acad. Sci. USA 109: 3434–3439.

Pasyanos, M. E. et al. 2014. LITHO1. 0: An updated crust and lithospheric model of the Earth. J. Geophys. Res., 119: 2153–2173.

Pellissier, L. et al. 2014. Quaternary coral reef refugia preserved fish diversity. – Science. 344: 1016– 1019.

Pulido-Santacruz, P. and Weir, J. T. 2016. Extinction as a driver of avian latitudinal diversity gradients. – Evolution 70: 860–872.

Pyron, R. A. 2014. Temperate extinction in squamate reptiles and the roots of latitudinal diversity gradients. – Glob. Ecol. Biogeogr. 23: 1126–1134.

Qian, H. et al. 2007. Environmental determinants of amphibian and reptile species richness in China. – Ecography 30: 471–482.

Raven, P. H., and Axelrod, D. I. 1975. History of the Flora and Fauna of Latin America: the theory of plate tectonics provides a basis for reinterpreting the origins and distribution of the biota. – American Scientist 63: 420–429

Renema, W. et al. 2008. Hopping hotspots: global shifts in marine biodiversity. – Science 321: 654– 657.

Richardson, J. E. et al. 2014. The influence of tectonics, sea-level changes and dispersal on migration and diversification of Isonandreae (Sapotaceae). – Bot. J. Linn. Soc. 174: 130–140.

Ricklefs, R. 2004. A comprehensive framework for global patterns in biodiversity. – Ecol. Lett. 7: 1– 15.

Ricklefs, R. et al. 2004. The region effect on mesoscale plant species richness between eastern Asia and eastern North America. – Ecography, 27: 129–136.

Ricklefs, R. E. and He, F. 2016. Region effects influence local tree species diversity. – Proc. Natl Acad. Sci. USA, 113: 674–679.

Rahbek, C. et al. 2007. Predicting continental-scale patterns of bird species richness with spatially explicit models. – Proc. R. Soc. Biol. Sci 274: 165–174.

Sandel, B. et al. 2011. The influence of late Quaternary climate-change velocity on species endemism. - Science 334: 660–664.

Sepulchre, P. 2006. Tectonic uplift and Eastern Africa aridification. – Science, 313: 1419–1423.

Seton, M. et al. 2012. Global continental and ocean basin reconstructions since 200Ma. Earth-Sci. Rev. 113: 212–270.

Sommer, B. et al. 2014. Trait-mediated environmental filtering drives assembly at biogeographic transition zones. – Ecology 95: 1000–1009.

Steeman, M. E. et al. 2009. Radiation of Extant Cetaceans Driven by Restructuring of the Oceans. – Syst. Biol. 58: 573–585.

Stein, A. 2014. Environmental heterogeneity as a universal driver of species richness across taxa, biomes and spatial scales. – Ecol. Lett. 17: 866–880.

Svenning, J. C. 2015. The influence of paleoclimate on present-day patterns in biodiversity and ecosystems. – Ann. Rev. Ecol. Evol. Syst. 46: 551–572.

Toussaint, E. F. 2014. The towering orogeny of New Guinea as a trigger for arthropod megadiversity. Nature Comm. 5.

Valentine, J. W., and E. M. Moores. 1970. Plate-tectonic regulation of faunal diversity and sea level: a model. – Nature 228: 657–659.

Valentine, J. W. (1971). Plate tetonics and shallow marine diversity and endemism, an actualistic model. – System. Biol. 20: 253–264.

Waide, R. B. 1999. The relationship between productivity and species richness. – Ann. Rev. Ecol. Evol. Syst. 30: 257–300.

Warren, B. H. et al. 2010. Why does the biota of the Madagascar region have such a strong Asiatic flavour? – Cladistics 26: 526–538.

Wilf, P. et al. 2005. Eocene plant diversity at Laguna del Hunco and Río Pichileufú, Patagonia, Argentina. – Am. Nat. 165, 634–650.

Xiang, Q. and Soltis, D. E. 2001. Dispersal-vicariance analyses of intercontinental disjuncts: historical biogeographical implications for angiosperms in the northern hemisphere. – Int. J. Plant Sci. 162: S29–S39.

